# Rapid assembly of functional modules for generating human artificial chromosome constructs compatible with epigenetic centromere seeding

**DOI:** 10.1101/2025.10.02.680129

**Authors:** Gabriel J. Birchak, Daniel G. Gibson, Praveen Kumar Allu, Prakriti Kashyap, John I. Glass, Ben E. Black

**Affiliations:** Department of Biochemistry & Biophysics, Perelman School of Medicine, University of Pennsylvania, Philadelphia, PA, 19104 USA; Penn Center for Genome Integrity, Perelman School of Medicine, University of Pennsylvania, Philadelphia, PA, 19104 USA; Epigenetics Institute, Perelman School of Medicine, University of Pennsylvania, Philadelphia, PA, 19104 USA; Graduate Program in Cell & Molecular Biology, Perelman School of Medicine, University of Pennsylvania, Philadelphia, PA, 19104 USA; J. Craig Venter Institute, La Jolla, CA 92037 USA

## Abstract

The ongoing development of human artificial chromosomes (HACs) will permit investigation into essential centromere processes and the means to deliver large genetic cargoes to target cells. Starting with large (∼750 kb) yeast artificial chromosome (YAC)-based constructs limits the rampant multimerization that has complicated many prior types of HACs. Large YAC construction is accomplished using transformation-associated recombination (TAR) strategies that can become unwieldly when several functional modules are to be incorporated and tested. To address this issue, we developed an approach where modules are built using high-fidelity *in vitro* assembly strategies in a bacterial artificial chromosome (BAC) format. Then, the assembled modules are transferred in a simplified TAR step into a recipient YAC harboring the prokaryotic “stuffer” DNA that comprises a large portion of the final HAC construct. This approach is highly efficient with two-thirds of all screened yeast clones harboring the correct TAR product. Further, whole-genome Oxford Nanopore Technologies (ONT) sequencing/alignments, *de novo* assembly of the final YAC using a single ONT sequencing run, and close inspection of highly repetitive regions are all streamlined to rapidly validate clones that match the design. The fully sequenced, verified strain harboring a multi-module construct was then fused to human cells, where it efficiently formed functional HACs upon initial seeding with CENP-A-containing nucleosomes. We envision that the rapid assembly steps will be useful to quickly incorporate different functional modules, including diverse genetic cargoes, to engineer HACs with specific design features.

## Introduction

Human artificial chromosomes (HACs) that form *de novo* in recipient human cells have typically been derived from DNA introduced by bacterial artificial chromosomes (BACs) or yeast artificial chromosomes (YACs) that then acquires the ability to replicate in S-phase and subsequently segregate like natural chromosomes to daughter cells at cell division (Harrington et al., 1997; Ikeno et al., 1998; Okada et al., 2007; Logsdon et al., 2019). Chromosome segregation is mediated by the centromere (Kixmoeller et al., 2020; Musacchio & Desai, 2017), which is defined by chromatin harboring the histone H3 variant, CENP-A (Earnshaw & Rothfield, 1985; Palmer et al., 1987). CENP-A nucleosomes recruit a 16-protein complex, the constitutive centromere-associated network (CCAN) (Amano et al., 2009; Foltz et al., 2006; Hori et al., 2008; Okada et al., 2006), that is present throughout the cell cycle and provides the foundation for the mitotic kinetochore that connects to spindle microtubules. Rare, spontaneous acquisition of functional centromeric chromatin can be used to identify DNA sequence features in human α- satellite DNA that contribute to natural chromosome behavior (Hayden et al., 2013; Maloney et al., 2012; Ohzeki et al., 2002; Schueler et al., 2001). Efficient centromere formation on α- satellite or even non-repetitive DNA lacking sequence homology to α-satellite DNA can be achieved by experimentally directing CENP-A nucleosome assembly (Barnhart et al., 2011; Chen et al., 2014; Hori et al., 2012; Mendiburo et al., 2011). The same approach has facilitated more recent efforts in the efficient formation of HACs (Logsdon et al., 2019).

The majority of *de novo* HAC formation approaches since their inception involved initial DNA constructs in the 40-200 kb range that invariably were rearranged (with themselves or with host cell DNA) and then multimerized many times to final sizes in the megabase range (Ebersole et al., 2000; Harrington et al., 1997; Ikeno et al., 1998; Kouprina et al., 2012; Logsdon et al., 2019; Rudd et al., 2003). These rearrangements and amplifications are not compatible with control over the genetic material carried by the HACs. Using a larger initial YAC construct (∼750 kb)—*sufficient for chromatin spanning from the kinetochore to the inner centromere where sister chromatin cohesion is maintained until the moment of their separation at the onset of anaphase* —along with epigenetic centromere seeding and an improved cell delivery strategy successfully leads to efficient HAC formation without rampant multimerization (Gambogi et al., 2024).

Outfitting the ∼750 kb YAC involved a cloning strategy that required the simultaneous recombination of several different DNA fragments in yeast by transformation-associated recombination (TAR) (Gambogi et al., 2024). Simultaneous assembly of multiple DNA fragments substantially reduces TAR efficiency (Gibson et al., 2008a; Gibson et al., 2008b), however, and is likely to do so to such an extent with some designs as to preclude isolation of the desired YAC.

Here, we report an approach that combines modules in high-efficiency steps that precede a TAR step to introduce designed HAC modules into a general YAC encoding the prokaryotic “stuffer” DNA. The initial steps involve rapid cycles of standard isothermal assembly, isolation in *E. coli*, and DNA sequence validation. After multiple modules are assembled in a BAC, it is introduced via a simplified high-efficiency TAR reaction. Further, the YAC built from component modules in this fashion efficiently forms functional HACs in recipient human cells by epigenetic centromere seeding.

## Results and Discussion

### Combining functional modules prior to TAR-mediated assembly of a HAC construct

We redesigned the strategy (Fig. 1) to assemble a functional HAC construct with the same basic features of one that we recently reported, termed YAC-*Mm*-4q21^LacO^ (Gambogi et al., 2024). As in the initial effort, the starting point is a YAC harboring the *M. mycoides* ∼550 kb synthetic prokaryotic genome (JCVI-syn3B; (Bittencourt et al., 2024)) that will serve as stuffer DNA so that the overall HAC construct can function without the need to increase size via multimerization or acquisition of genomic DNA (Gambogi et al., 2024; Logsdon et al., 2019). The HAC construct design requires the addition to the starting YAC of four functional modules: 1) a fluorescent protein to validate spheroplast fusion to human cells; 2) an antibiotic resistance gene to select for cells that harbor the HAC construct; 3) an array of bacterial Lac operator (LacO) sites for initial local assembly of CENP-A nucleosomes via a pulse of expression in recipient cells of a fusion protein of the CENP-A-specific histone chaperone, HJURP, and the Lac repressor (LacI); and 4) a 182 kb human chromosome segment (4q21) within which functional centromeric chromatin can form (Logsdon et al., 2019). A fifth module, a yeast auxotrophic selection marker, is also needed to select for yeast candidates that performed TAR.

**Figure 1.**
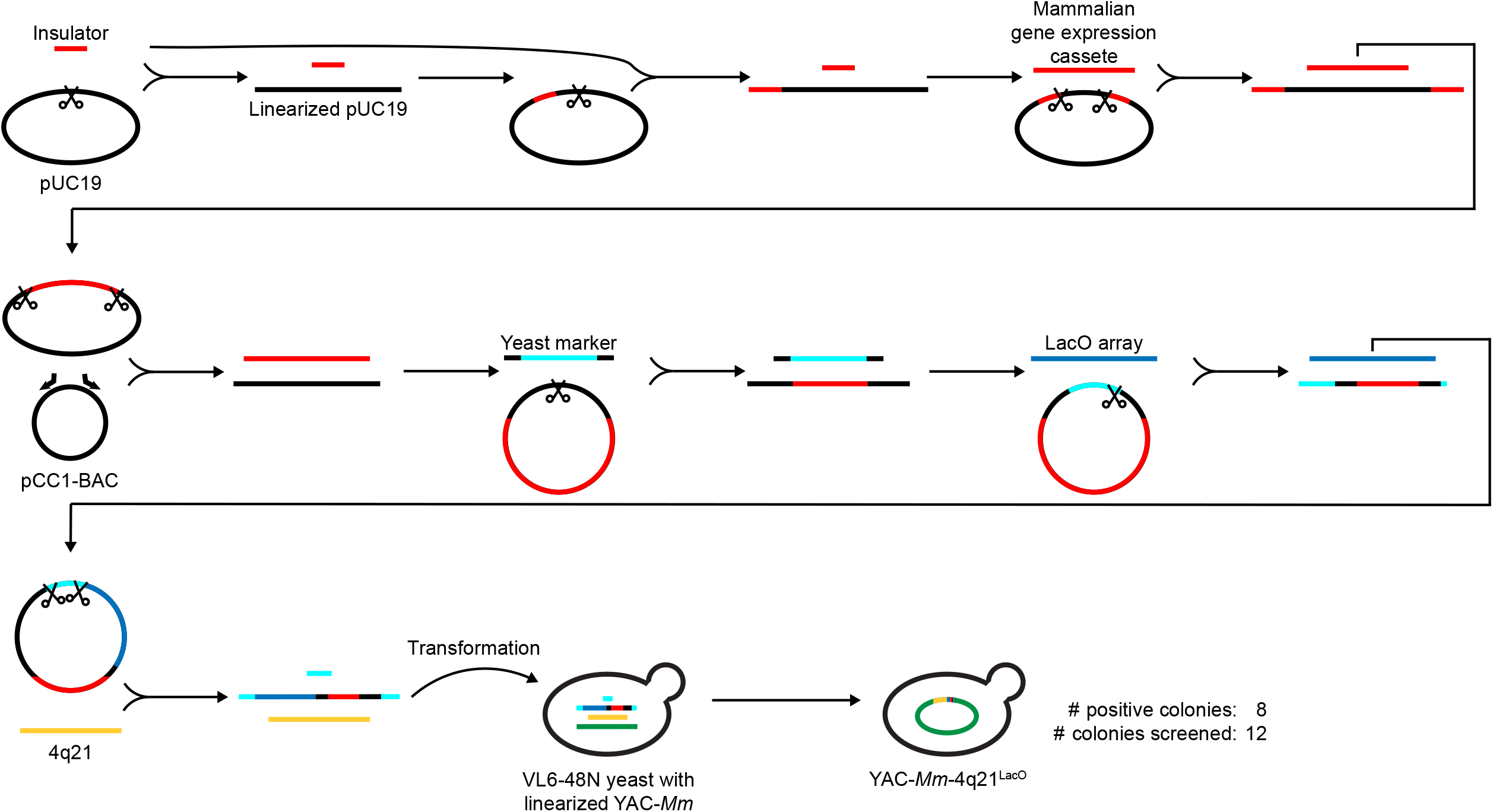
Synthesizing a multi-module BAC for simplified and efficient TAR cloning to build a new YAC-*Mm*-4q21^LacO^. Graphical depiction of cloning strategy to generate YAC-*Mm*-4q21^LacO^ in a single TAR step. First, a pair of tRNA dimers serving as insulators were inserted into the pUC19 vector. Subsequently, this vector was digested between the insulators and ligated to the *hEF1α>tdTomato-P2A-NeoR* expression cassette. Next, this recombinant plasmid was digested, and the insulator-flanked gene expression cassette was transferred to the linearized pCC1-BAC by isothermal assembly. This recombinant BAC was digested and mixed with a landing pad containing an auxotrophic marker, an 80-bp YAC hook with an internal restriction site, 40 bp of homology to the vector backbone housing the neocentromere-adjacent 4q21 locus, and 40 bp of homology to the vector backbone housing a 10 kb LacO array. Next, the vector was digested within the LacO landing pad and then isothermally assembled with the linearized LacO array, resulting in a BAC carrying both the LacO array and the gene expression cassette. In the final step, the BAC was digested in the YAC hook and in the 4q21 landing pad and co-transformed with the linearized 4q21 BAC and an sgRNA into YAC-*Mm* yeast constitutively expressing Cas9 to carry out TAR, generating the final YAC-*Mm*-4q21^LacO^ construct. Scissor symbols indicate restriction digestions at the corresponding cloning steps. Bent arrows indicate PCR primers sites for the corresponding cloning steps. Note that DNA fragments and vectors are not shown to scale for the sake of simplicity in diagramming out the overall strategy.

Each of the intermediate steps in the assembly are built into standard plasmids (e.g. pUC19) or BACs (e.g. pCC1) (Fig. 1). The inserts include the synthesized mammalian expression cassette encoding both the fluorescent (tdTomato) and antibiotic (neomycin)-resistance proteins flanked by tRNA dimers (Ebersole et al., 2011) that are known to resist silencing mediated by pericentric heterochromatin (Blower & Karpen, 2001; He & Brinkley, 1996; Lee et al., 2013; Partridge et al., 2000), PCR amplicon (*C. albicans* URA3 gene), and digestion product from an existing vector (including the ∼10 kb LacO array (Robinett et al., 1996)). At each step, homology “hooks” of ∼40 bp were deployed for use with well-established and highly efficient isothermal assembly cloning methodologies (Gibson et al., 2009). Indeed, all cloning steps through the final BAC prior to TAR cloning can be realistically accomplished by a single researcher in less than 3 weeks.

To expedite YAC assembly, this final BAC containing the engineered modules (with 4q21 hooks) and the BAC carrying 4q21 were linearized and co-transformed with an sgRNA into YAC-*Mm* yeast constitutively expressing Cas9 (Kannan et al., 2016) to carry out TAR, generating the final YAC-*Mm*-4q21^LacO^ construct. As expected with the relatively simple TAR reaction, hundreds of colonies survived auxotrophic selection. Further, 8/12 colonies screened by junctional PCR yielded products indicative of proper TAR assembly (Fig. 1). These results provided an early indication that our rapid HAC construct assembly strategy achieves the objective of efficiency and accuracy.

### Genomic sequencing of the strain harboring the newly built YAC-*Mm*-4q21^LacO^

We performed long-read ONT whole-genome sequencing of a candidate yeast strain that had passed PCR-based diagnostic screening and acquired sufficient coverage to perform an unbiased whole-genome assembly. Alignment of reads back to the assembled YAC showed contiguous coverage (Fig. 2A). Alignment also reveals lower depth of coverage of the prokaryotic stuffer DNA (Fig. 2A). Given that the depth of coverage of other portions of the construct is comparable to that of the native yeast chromosomes (Fig. S1A), we concluded that lower depth of coverage in the stuffer DNA is due to an underrepresentation of many of the prokaryotic reads, possibly related to a technical bias against its extremely high A/T content. In support of this conclusion, an independently isolated TAR candidate exhibited a similar underrepresentation of much of the prokaryotic portions of the construct (Fig. S1B). The sequencing also showed that the second sequenced candidate is essentially identical to our initial candidate TAR-generated YAC, except for of it having fewer LacO copies. Further analysis of the initial candidate YAC involved closer inspection of reads that span each junction, confirming seamless, on-target integration (Fig. 2B,C). We also leveraged the long-read sequencing modality to unambiguously evaluate LacO array length and stability by selectively examining reads that spanned the entire array (i.e., had homology to unique sequence flanking the array). In doing so, we found a median LacO number of 259 among all reads with a large fraction (42/44) deviating by less than 10% (Fig. 2D), suggesting that the array is stable in culture despite the reported recombination proficiency of the yeast strain (Kouprina et al., 1998). Further, the genomic sequencing data was sufficient to assemble a circular 756 kb YAC that was mapped back to the designed reference sequence, confirming the absence of gross indels or rearrangements (Fig. 2E). This approach reveals a matrix associated with the human 4q21 region, a feature in our graph that we expected due to Alu elements that are highly abundant in 4q21 and throughout the human genome (Houck et al., 1979). The long-read sequencing data was also of sufficient quality to unambiguously assemble all native yeast chromosomes (Fig. 2F). Together, our analysis of the strain harboring YAC-*Mm*-4q21^LacO^, revealed the desired TAR-mediated assembly alongside an intact, natural yeast genome.

**Figure 2.**
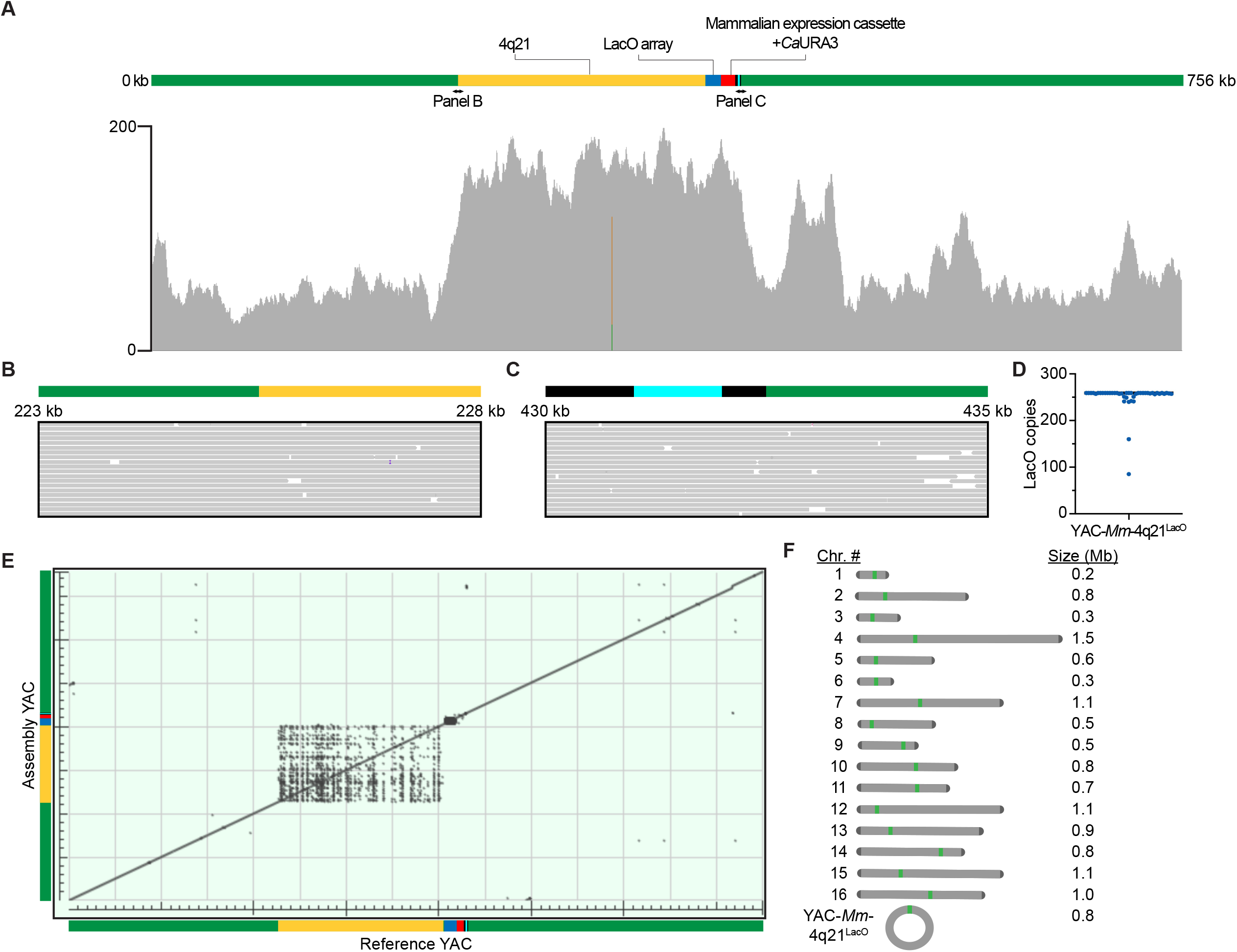
Long-read DNA sequencing of the yeast genome harboring YAC-*Mm*-4q21^LacO^. A) Integrated Genomics Viewer alignment track showing raw reads mapped to the *de novo* assembled YAC-*Mm*-4q21^LacO^. Grey denotes a perfect match, copper denotes the unexpected presence of a guanine, and green denotes the unexpected presence of an adenine. In total, these unexpected instances impact 1 bp out of the entire 756 kb construct. B) Reads spanning a 5 kb region that includes the upstream junction of the BAC integration site. The purple bar represents a nucleotide insertion. Insertions that do not comprise consensus are a result of the error rate inherent to sequencing. C) Reads spanning a 5 kb region that includes the downstream junction of the BAC integration site. D) Distribution of LacO copies for each read spanning the entire array. Bar denotes median LacO copy number of 259 for the 44 reads (individually represented by blue dots) analyzed. E) Dot plot generated by BLAST alignment of the *de novo* assembled YAC-*Mm*-4q21^LacO^ against the predicted recombinant YAC, referred to as “Reference YAC.” F) Graphical depiction of the YAC-*Mm*-4q21^LacO^ genome. All chromosomes except native chromosome 12 and lengths are derived from unbiased, genome assembly. *De novo* assembly failed to resolve the rDNA locus on chromosome 12, resulting in two contigs as previously reported (Salazar et al., 2017; Yue et al., 2017). Consequently, budding yeast chromosome 12 was manually assembled by joining the two contigs. Green denotes the yeast centromere, and dark grey denotes yeast telomeres.

### HAC formation with YAC-*Mm*-4q21^LacO^ by spheroplast fusion and centromere seeding

We tested the ability of our newly built YAC-*Mm*-4q21^LacO^ construct to form HACs (Fig. 3) using the same general strategy (Fig. 3A) as reported with the prior version that had been generated through an independent cloning strategy (Gambogi et al., 2024). This involved YAC- *Mm*-4q21^LacO^ delivery to HT1080^Dox-inducible mCherry-LacI-HJURP^ or U2OS^Dox-inducible mCherry-LacI-HJURP^ cells via spheroplast fusion. An initial 48 hr pulse of doxycycline (dox) was then applied to recipient cells before subjecting the cells to G418 selection. Surviving cells were expanded to screen for HAC formation events by a combined florescence *in situ* hybridization (using probes targeting the *M. mycoides* stuffer that comprises ∼550 kb of YAC-*Mm*-4q21^LacO^) and immunofluorescence to detect human CENP-A (Fig. 3A). As we anticipated based upon the earlier version of this YAC (Gambogi et al., 2024), formation of HACs was stimulated by the early pulse of dox-inducible mCherry-LacI-HJURP expression in either HT1080 or U2OS backgrounds (Fig. 3B,C). Further, we note that the detected HACs (Fig. 3) were not of the large size associated with multimerization from smaller BACs, as in prior approaches with smaller BAC-based HAC formation approaches (Logsdon et al., 2019). Rather, they are small, consistent with our initial version of YAC-*Mm*-4q21^LacO^ that avoids rampant multimerization during HAC formation (Gambogi, et al., 2024). Taken together, our HAC formation tests (Fig. 3) indicate efficient HAC formation with the newly constructed version of YAC-*Mm*-4q21^LacO^ is stimulated by epigenetic centromere seeding, in a similar manner as the original version (Gambogi et al., 2024).

**Figure 3.**
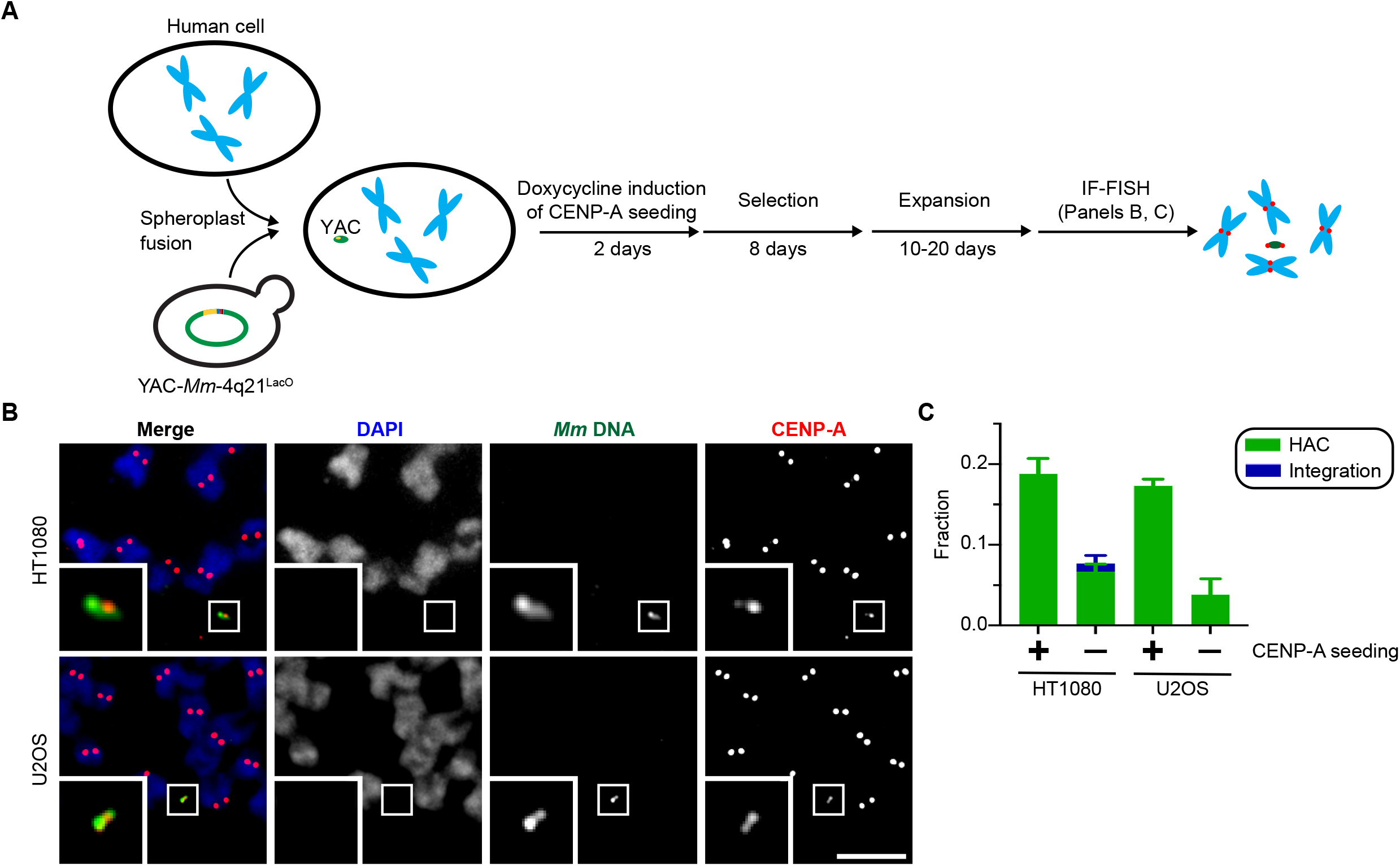
YAC-*Mm*-4q21^LacO^ built via simplified TAR cloning forms functional HACs upon centromere seeding in human cells. A) Graphical representation of the HAC formation assay. B) Representative HACs formed after an initial pulse of mCherry-LacI-HJURP expression in HT1080 and U2OS cells. Inset, 5X magnification. Bar, 5 µm. C) Quantification of HAC formation frequency with and without doxycycline-mediated induction. The mean (+/- standard error) is shown. The increase in HAC frequency was assessed using an unpaired, one-tailed T test. P(HT1080 + dox = HT1080 - dox) < 0.001, n=4 (≥ 20 cells, replicate). P(U2OS + dox = U2OS – dox) < 0.01, n=3 (≥ 20 cells, replicate).

## Conclusions

HACs are attractive for their ability to test designed functional modules. YAC-based HAC construct assembly coupled with direct delivery to target human cells via spheroplast fusion promises to expand these modules in terms of their number and size due to their large carrying capacity. This promise raises new challenges, including the ability to construct YACs of increasing complexity. In this study, we address one of these new challenges: the need for rapid and efficient HAC construct assembly. Without addressing this need, some TAR reactions are so inefficient that it is easy to imagine rounds of TAR attempts yielding no proper assemblies (i.e., no colonies surviving auxotrophic selection or no surviving colonies having correctly assembled recombinant YACs). What is needed is an approach that can be standardized to faithfully engineer HAC-precursor YACs. We achieved this with the new assembly of YAC-*Mm*-4q21^LacO^ by moving the majority of steps (Fig. 1) to *in vitro* assembly methodologies already proven to be rapid and efficient (Gibson et al., 2009). We note that the modules include potentially challenging DNA sequences, both in terms of size (which could hamper biochemical isolation and transformation into yeast, e.g., the 182 kb human chromosome 4q21 sequence) and repetitiveness (which could complicate the necessary recombination step in TAR cloning, e.g., the LacO array). The pipeline to couple YAC construction analysis to contemporary long-read sequencing methodologies is highly effective and is rendered rapid and efficient by well-established in-house and commercial microbial whole genome sequencing platforms. In sum, this advance is an important one among many that will enhance access to the design, synthesis, and testing of new HACs that address goals of diverse human chromosome engineering efforts.

## Methods

### BAC cloning

Four Integrated DNA Technologies (IDT) Ultramers (5’- TAGTGGTGAGCATAGCTGCCTTCCAAGCAGTTGACCCGGGTTCGATTCCCGGCCAACGCAGCGTGCCCACGTTTTGCCTGGCGCCGCGTGCTAGCAAAGGTCCCTTTTCACAGAGCAGTTTTGAAACACTGTTTTTGTAGGTTTTCCAAGGGGATAATTATAGCTCATTGAGCTTTGGCGCGTGCTCCCTGGTGGTCTAG-3’, 5’- CCAGCAGAGAGGAAGAAGAGAGGCTTCCCTGACCGGGAATCGAACCCGGGCCGCGGCGGTGAGAGCGCCGAATCCTAACCACTAGACCACCAGGGAGCACGCGC-3’, 5’- ACGCGCCAATCCCATTGCAAATTCTACAAAAGGAGTGTTTCCCAACTGCTCTATCAAGAGGAATGTTGCACACTGTGACCTGAATGCAAACATCACACGCGCCAGCAGAGAGGAAGAAGAGAGGCTTCCC-3’, 5’- ACGCGTGTGATGGCTGCATTCCACACACAAGGTGGAACATTTCTCTTGATAGAGCAGTTTTGAAACACTCTTCCCGTAGAATCTGCTGGCGCGTGTGCGCGTTGGTGGTATAGTGGTGAGCATAGCTGCCTTCCAAGC-3’) were used in an overlap extension PCR reaction (primers: 5’-ACGCGCCAATCCCATT-3’ and 5’-ACGCGTGTGATGGCT-3’) to generate a dimer of tRNA^Gly^ and tRNA^Glu^ genes (Ebersole et al., 2011). These dimers were serially incorporated into a plasmid by isothermal assembly, resulting in a vector with an inverted dimer pair.

Isothermal assembly was performed using NEB HiFi 2X master mix (NEB catalog #: E2621) according to the manufacturer’s instructions. This vector and the plasmid carrying *hEF1α*>*tdTomato-P2A-NeoR-SV40*^*polyA*^ (synthesized by Twist Biosciences), were digested with enzymes SphI-HF and SalI-HF. The vector backbone including the insulators and the expression cassette insert were gel extracted using the QIAquick Gel Extraction Kit (Qiagen catalog #: 28704) and used in a two-fragment ligation with T4 DNA ligase (NEB catalog #: M0202) according to manufacturer instructions with the ligation being carried out at 16 °C overnight. The plasmid carrying the insulator-flanked gene expression cassette was validated by Plasmidsaurus sequencing.

The plasmid was digested by PvuII-HF to release the insulator-flanked expression cassette and isothermally assembled to AfeI-digested pCC1-BAC vector, which was PCR amplified at that site to contain 40 bp homology arms to the PvuII-released cassette. In a second round of DNA assembly, this new plasmid was digested with NotI and combined with the modular landing pad consisting of *C. albicans URA3* (PCR amplified from pAG60 plasmid (Addgene) using primers 5’-ATAGGCCACTAGTGGATCTGATATCATCGATGAATTCGAG-3’ and 5’-CTGTTTAGCTTGCCTCGTCC-3’), two 40-bp hooks for the microbial stuffer, two 40- bp hooks for 4q21 incorporation, and two 40-bp hooks for LacO array incorporation. This vector was then digested with AscI, and the plasmid carrying the LacO array (Logsdon et al., 2019) was digested with AseI and MfeI before combining these two fragments in a third round of isothermal assembly.

TAR cloning to generate YAC-*Mm*-4q21^LacO^ yeast For TAR, the LacO-containing BAC was digested with PacI and NotI, and the BAC carrying the 4q21 locus (Logsdon et al., 2019) was digested with NotI. Both digests were phenol-chloroform extracted and mixed with an sgRNA generated by IDT (5’- mGmAmAGAUAAAAAUCCUCUAUCGUUUUAGAGCUAGAAAUAGCAAGUUAAAAUA AGGCUAGUCCGUUAUCAACUUGAAAAAGUGGCACCGAGUCGGUGCmUmUmUU-3’) to target the Syn3B stuffer constituting YAC-*Mm*. Spheroplast transformation was carried out as follows. YAC-*Mm* yeast (Bittencourt et al., 2024) (VL6-48N strain carrying YAC-*Mm* and constitutively expressing Cas9 (Kannan et al., 2016)) were grown in 50 ml synthetic minimal media -histidine to an OD_600_ of 1.0. The culture was pelleted at 1,600 x g for 3 min, washed with 50 ml sterile water, pelleted again, and resuspended in 20 ml 1 M sorbitol. The yeast were incubated on ice at 4 °C overnight. The next day, cells were harvested at 1,741 x g for 3 min and resuspended in 20 ml SPE (1 M sorbitol, 7.89 mM Na_2_HPO_4_, 2.33 mM NaH_2_PO_4_, 10 mM EDTA pH 8.0). 40 µl of β-mercaptoethanol and 40 µl 10 mg/ml zymolyase-20T were added to the suspension. The tube was inverted several times and then placed in a shaker to incubate at 30 °C, 50 rpm. After 20 min, the tube was inverted several times and again returned to the shaker to incubate for 20 min. 30 ml of 1 M sorbitol was added to the suspension before harvesting spheroplasts at 1,600 x g for 5 min. Spheroplasts were washed with 50 ml 1 M sorbitol before harvesting at 1,600 x g for 5 min. Spheroplasts were resuspended in 2.5 ml STC (1 M sorbitol, 10 mM Tris-Cl pH 7.5, 10 mM CaCl_2_) and incubated at room temperature for 10 min. 200 µl of the suspension was added to an Eppendorf tube containing the DNA and sgRNA along with water to a total volume of 40 µl, and the suspension was incubated for 10 min at room temperature. 1 ml of PEG/CaCl_2_ (20% w/v PEG8000, 10 mM CaCl_2_, 10 mM Tris-Cl pH 7.5) was added and the solution mixed by inverting and flicking the Eppendorf tube. The solution was incubated at room temperature for 20 min. Transformed spheroplasts were harvested at 1,600 x g for 8 min. The spheroplasts were suspended in 800 µl SOS medium (1 M sorbitol, 6.5 mM CaCl2, 0.25% w/v bacto yeast extract, 0.5% w/v bacto peptone) and incubated at 30 °C for 1 h.

The 800 µl solution was then pipetted onto 8 ml of 55 °C synthetic minimal top agar, inverted 3 times, and poured onto synthetic minimal sorbitol plates for recovery of transformants. -Ura prototrophs were patched onto synthetic minimal -histidine -uracil plates for genotyping.

Patches were screened by direct PCR without DNA extraction. Briefly, a QIAGEN Multiplex PCR Kit (cat # 206145) was used to prepare a PCR master mix containing primers for either the left junction or the right junction. A small amount of cells from each of the patches were combined with 15 µl of the master mix and cycled per the instructions.

YAC-*Mm*-4q21^LacO^ genome assembly and analysis YAC-*Mm*-4q21^LacO^ yeast were grown in 50 ml of synthetic minimal media (-histidine and -uracil) to OD_600_=1.0. Yeast cells were pelleted at 500 x g for 2 min. The pellet was resuspended in .5 ml of DNA/RNA shield (Zymo Research catalog #: R1100) before sending to Plasmidsaurus for ONT sequencing.

Raw reads from the ONT run were assembled using Flye (v. 2.9.2) in nano-hq mode while specifying the genome length as 13 Mb. Reads were mapped back to this *de novo* assembly using Minimap2 (v. 2.28-r1209), ensuring assembly contiguity. Samtools (v. 1.6) was used to filter out secondary and supplementary alignments, sort remaining primary aligned reads, and index aligned reads. The resulting alignment was visualized in the Integrated Genomics Viewer (v. 2.13.2).

To evaluate LacO array length, Seqtk (v. 1.4-r122) was used to retrieve pass-filter reads mapping to 300 unique nucleotides upstream and downstream of the LacO array. The length of each LacO array, considered the distance between the first bp of the first monomer and the last bp of the last monomer, was divided by the number of bp per monomer (39.85, including the monomer and intervening sequence) to generate the number of LacO copies for each read.

### Spheroplast fusion

YAC-*Mm*-4q21^LacO^ yeast were grown in synthetic minimal media (-histidine -uracil) to OD_600_ = 0.8-1.0 before harvesting at 1,741 x g for 3 min. Yeast were resuspended in 20 ml 1 M sorbitol and incubated overnight at 4 °C. Yeast were harvested at 1,741 x g for 3 min and resuspended in 20 ml SPE medium. 40 µl β-mercaptoethanol and 100 µl 10mg/ml zymolyase-20T were added to the yeast solution before gently inverting and then incubating at 37 °C in a shaker set to 25 rpm for 1 h. 30 ml ice cold 1 M sorbitol was then added to the spheroplasts before pelleting at 627 x g, 18 min, 4 °C. The spheroplasts were washed in 50 ml ice cold 1 M sorbitol before harvesting at 627 x g, 25 min, 4 °C. The pellet was resuspended in 1 ml STC, and the solution was allowed to incubate at room temperature for 10-30 min before addition to the HT1080^Dox-inducible mCherry-LacI-HJURP^ cells or U2OS^Dox-inducible mCherry-LacI-HJURP^. The cells were then processed along with the spheroplasts. ∼4 h before yeast were pelleted, HT1080^Dox-inducible mCherry-LacI-HJURP^ cells or U2OS^Dox-inducible mCherry-LacI-HJURP^ cells (∼60-80% confluent) were supplied with fresh DMEM containing 10% certified tetracycline-free fetal bovine serum (Takara catalog #: 631106), 100 U/ml penicillin, and 100 µg/ml streptomycin (hereafter referred to as complete DMEM) and 50 µM S-trityl-L-cysteine (STLC) (Fisher: AAL1438406). After 5-6 h incubation in 50 µM STLC, cells were washed with PBS, trypsinized, and counted using a hemacytometer.

Cells were pelleted and resuspended in PBS to a concentration of 600,000 cells/ml. 300,000 HT1080^Dox-inducible mCherry-LacI-HJURP^ cells or U2OS^Dox-inducible mCherry-LacI-HJURP^ cells were mixed with 90,000,000 YAC-*Mm*-4q21^LacO^ spheroplasts and incubated at room temperature for 5 min. The mixture was pelleted at 1,700 x g for 30 s in a microcentrifuge. The supernatant was aspirated and the pellet resuspended in a 45% PEG2000, 10% DMSO solution in 75 mM HEPES pH 7.5. The suspension was incubated at room temperature for 5 min. The fusion was quenched by adding 1 ml serum/antibiotic-free DMEM. Fused cells were pelleted at 1,700 x g for 30 s in a microcentrifuge. Cells were resuspended in 1 ml complete DMEM and 500 µl aliquoted into a 6-well plate containing 2 ml complete DMEM. After 4 h (when cells have adhered), media is swapped with 3 ml complete DMEM +/- 2 µg/ml doxycycline. After 16 h, cells are again supplied with 3 ml complete DMEM +/- 2 µg/ml doxycycline. Cells are allowed to incubate for 32 h before washing with PBS, trypsinizing, and transferring to a 10-cm plate with complete DMEM + 325 µg/ml G418. Selection is maintained for 8 days before reducing selection to the 150 µg/ml G418 maintenance concentration for subsequent expansion.

### IF-FISH on metaphase spreads

70-80% confluent human cells were fed with complete DMEM containing 50 µM STLC and incubated at 37 °C for 2-4 h. Mitotic cells were blown off using a transfer pipet and pelleted at 415 x g for 5 min. Cells were resuspended in 500 µl PBS before counting with a hemacytometer. The cells were again pelleted at 415 x g for 5 min before dropwise resuspending the cells in hypotonic buffer consisting of a 1:1:1 ratio of 75 mM KCl, 0.8% Na_3_C_6_H_5_O_7_, and a solution consisting of 1.5 mM MgCl_2_ and 3 mM CaCl_2_ to a concentration of 500,000 cells/ml.

Cells were allowed to swell at room temperature for 15 min before diluting to 50,000 cells/ml in 500 µl total hypotonic buffer. Cells were cytospun at 1500 rpm, acceleration 9, for 5 min in a Shandon Cytospin 4 centrifuge. Cells were allowed to dry for 1 min before permeabilizing with KCM buffer (10 mM Tris-Cl pH 7.7, 120 mM KCl, 20 mM NaCl, and 0.1% triton X-100) for 15 min. Spreads were blocked in IF block (2% FBS, 2% BSA, 0.1% tween-20, and 0.02% NaN_3_) for 20 min before incubating with αCENP-A mAb (ENZO catalog #: ADI-KAM-CC006) diluted to 1 µg/ml in IF block for 45 min at room temperature. Spreads were washed 3 times in KCM for 5 min each before incubating in Cy3-conjugated donkey anti-mouse IgG (Jackson Immunoresearch catalog #: 712-165-151) diluted to 3.5 µg/ml in IF block for 25 min. Spreads were again washed 3 times in KCM for 5 min before fixing in 4% paraformaldehyde for 10 min at room temperature. Spreads were washed 3 times with dH_2_O before proceeding with FISH.

FISH probe was prepared by nick translation (Roche catalog #: 10976776001) using biotin-16-dUTP (Millipore catalog #: 11093070910) and *M. mycoides* gDNA as template. Unincorporated nucleotides were removed using Amersham MicroSpin G-50 columns (Cytiva catalog #: 27533002). 300 ng of probe/slide was ethanol precipitated along with 20 µg of salmon sperm DNA (Invitrogen catalog #: 15632011) and 10 µg of human Cot1 DNA (Invitrogen catalog #: 15279011). Precipitated DNA was resuspended in 50% formamide, 10% dextran sulfate in 2X SSC and denatured at 78 °C for 10 min before being placed in 37 °C for at least 20 min for pre-hybridization. 300 ng of probe was added to the RNase-treated spreads, and hybridization was performed at 37 °C for 16 h. Slides were washed 2 times in 50% formamide, 2X SSC at 37 °C for 5 min before washing slides 2 times in 2X SSC at 37 °C for 5 min. Slides were blocked with 2.5% milk in 4X SSC + 0.1% tween-20 for 10 min and then incubated in NeutrAvidin-FITC (ThermoFisher catalog #: 31006) diluted to 25 µg/ml with 2.5% milk in 4X SSC + 0.1% tween-20 at 37 °C for 1 h. Spreads were washed 3 times in 4X SSC + 0.1% tween-20 at 45 °C at 45 °C for 2 min, followed by one wash in PBS. Spreads were then incubated in PBS supplemented with 1 µg/ml DAPI for 10 min at room temperature, washed again in PBS, and then water before mounting with Vectashield. Slides were imaged on an inverted epifluorescence microscope (Leica DMI6000B) with a 100x oil-immersion objective and the Leica K8 Scientific CMOS camera. Images were blind deconvolved within LAS X (v. 3.6.0.20104) while resizing from 12-bit depth to 16-bit depth. Channel contrasting and image cropping was carried out in ImageJ (v. 1.52k) before assembling figues in Adobe Illustrator (v. 29.7.1). The “HAC” designation was given to objects with FISH signal < 2 µm in diameter and two co-localized CENP-A foci. The “integration” designation was given to events with FISH signal located at the same site on each sister chromatid of a native chromosome and distal from the centromere.

## Acknowledgements

We thank our UPenn colleagues G.A. Logsdon and S-C. Chuang for advice related to ONT sequencing and analysis workflows and G.A. Logsdon for comments on the manuscript. The content is solely the responsibility of the authors and does not necessarily represent the official views of the National Institutes of Health. This manuscript is the result of funding in whole by the National Institutes of Health (NIH). It is subject to the NIH Public Access Policy. Through acceptance of this federal funding, NIH has been given a right to make this manuscript publicly available in PubMed Central upon the Official Date of Publication, as defined by NIH.

## Funding

Supported by NIH grants HG012445 (J.I.G. and B.E.B.) and CA265794 (B.E.B.). G.J.B. was supported by the Penn Cell and Molecular Biology Training Program: GM007229.

## Author Contributions

G.J.B., D.G.G., and B.E.B. conceived the study. G.J.B., D.G.G., P.K.A., and P.K. performed the experiments. G.J.B., P.K.A., P.K, and B.E.B. analyzed data. G.J.B. and B.E.B. wrote the paper. All authors edited the paper. J.I.G. and B.E.B. directed the research.

## Competing Interests

G.J.B., J.I.G., and B.E.B. are inventors on a patent application submitted by the University of Pennsylvania related to this work.

## Data and Materials Availability

Sequencing data will be released on the Sequence Read Archive (SRA) at the time of publication (accession #: PRJNA1322447).

---

**Figure S1.**
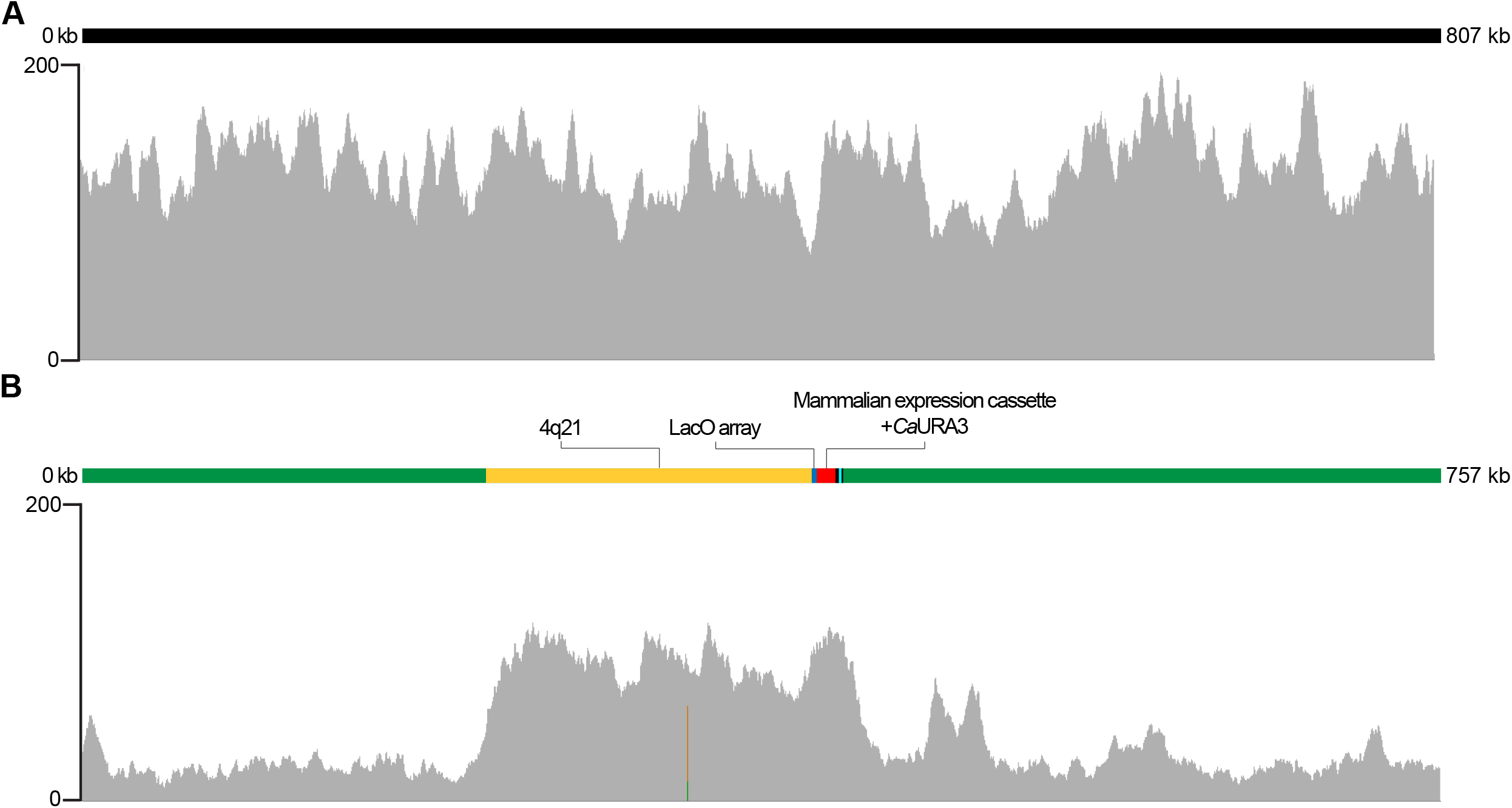
*M. mycoides* stuffer DNA is under-represented in sequencing libraries of recombinant yeast strains. A) An example Integrated Genomics Viewer alignment track showing coverage throughout the *de novo*-assembled native chromosome 2. B) Integrated Genomics Viewer alignment track showing raw reads mapped to *de novo*-assembled YAC-*Mm*-4q21^Short LacO^ again reveals relative underrepresentation of prokaryotic stuffer DNA.

